# A genome-wide CRISPR screen identifies UFMylation and TRAMP-like complexes as host factors required for hepatitis A virus infection

**DOI:** 10.1101/2019.12.15.877100

**Authors:** Jessie Kulsuptrakul, Ruofan Wang, Nathan L. Meyers, Melanie Ott, Andreas S. Puschnik

## Abstract

Hepatitis A virus (HAV) is a positive-sense RNA virus causing acute inflammation of the liver. Here, using a genome-scale CRISPR screen in a human hepatocyte cell line, we provide a comprehensive picture of the cellular factors, which are exploited by HAV during replication. We identified genes involved in sialic acid biosynthesis and members of the eukaryotic translation initiation factor complex, corroborating their putative roles in HAV infection. Additionally, we uncovered all components of the cellular machinery for UFMylation, a ubiquitin-like protein modification. We showed that HAV translation specifically depends on UFM1 conjugation of the ribosomal protein RPL26. Furthermore, we found that components related to the yeast Trf4/5– Air1/2–Mtr4 polyadenylation (TRAMP) complex, are required for viral translation, independent of controlling HAV poly(A) tails. While the identified HAV host factors were largely distinct compared to other picornaviruses, we highlighted a surprising co-dependency of HAV and hepatitis B virus (HBV) on the TRAMP-like complex. Finally, we demonstrated that pharmacological inhibition of the TRAMP-like complex decreased HAV replication in hepatocyte cells and human liver organoids, thus providing a strategy for host-directed therapy of HAV infection.

## Introduction

Hepatitis A virus (HAV) is a single-stranded, positive-sense RNA virus and a hepatotropic member of the *Picornaviridae*. Despite successes in vaccine development, HAV remains a common cause of enterically transmitted hepatitis globally, responsible for epidemics in developing and developed countries, and with symptoms ranging from mild inflammation and jaundice to acute liver failure (CDC, 2019; Jacobsen, 2018). However, no specific antiviral treatment is currently available. Thus, a better understanding of the molecular virus-host interactions is important and can provide insights for host-directed therapies.

During infection HAV enters host cells through interaction with cell surface molecules, followed by uncoating and delivery of viral RNA from endosomes to the cytosol. The genomic RNA is translated into a polyprotein and subsequently processed into the individual structural and non-structural proteins. HAV then co-opts cellular membranes to form virus-specific organelles, where genome replication occurs. Finally, the newly synthesized genomes are packaged and HAV egresses in the form of quasi-enveloped virions (McKnight and Lemon, 2018).

Despite a general understanding of the viral life cycle and suggested roles for certain cellular proteins during infection, a comprehensive assessment and thorough validation of the most critical host factors for HAV infection is lacking. For example, TIM1 (T-cell immunoglobulin mucin receptor 1), also named HAV cellular receptor 1 (HAVCR1), was originally identified as receptor for HAV but a more recent study disproved its essentiality for HAV infection (Das et al., 2017; Kaplan et al., 1996).

Additionally, numerous host proteins have been shown to interact with the viral RNA to potentially modulate translation or replication of the HAV genome. For example, the HAV internal ribosome binding site (IRES) is unique in that it requires an intact cellular eukaryotic translation initiation factor complex, while other picornaviruses shut down host mRNA translation by proteolytic cleavage (Ali et al., 2001; Borman and Kean, 1997). Other RNA binding proteins, such as PCBP2 (Poly(RC) binding protein 2), PTBP1 (Polypyrimidine tract-binding protein 1) or La protein have been shown to bind viral RNA, but their importance in viral translation or replication are not fully determined (Gosert et al., 2000; Graff et al., 1998; Jiang et al., 2014).

To address these knowledge gaps, we performed a genome-wide CRISPR knockout (KO) screen to identify host cellular factors that are critical for the HAV life cycle in an unbiased fashion. CRISPR screens have successfully been used to study human pathogens, such as mosquito-borne flaviviruses, enteroviruses or hepatitis C virus (HCV) (Diep et al., 2019; Marceau et al., 2016; Zhang et al., 2016).

Using this approach, we identified sialic acid and ganglioside biosynthesis genes, important for viral entry, as well as factors of the eukaryotic translation initiation factor complex and PTBP1, previously implicated in HAV translation. Additionally, we uncovered unknown cellular pathways, in particular UFMylation and a TRAMP-like complex (related to the yeast Trf4/5–Air1/2–Mtr4 polyadenylation complex), and pinpoint their role to HAV protein synthesis. Importantly, we demonstrate that these host factor requirements are unique to HAV compared to other picornaviruses. Surprisingly, we reveal that hepatitis B virus (HBV), a DNA virus with a vastly different replication strategy, also strongly depends on ZCCHC14, thus highlighting an unexpected convergence across different viral families to co-opt the same cellular complex. Finally, we provide proof-of-concept for host-directed therapy of HAV infections using a small molecule inhibitor against subunits of the TRAMP-like complex.

## Results

### A genome-wide CRISPR screen reveals novel host factors required for HAV infection

To identify host cellular factors that are critical for HAV infection, we conducted a genome-wide CRISPR KO screen in the human hepatocyte cell line Huh7.5.1 using the cytopathic HAV HM175/18f strain (Lemon et al., 1991) (Figure 1A). Infection led to notable cell death after 3 days and surviving cells were collected after 12 days for analysis. We found enrichment of guide RNA (gRNA) sequences targeting genes involved in several pathways known to be required for HAV replication (Figure 1B and Table S1). For example, sialic acid and ganglioside biosynthesis genes (*GNE, CMAS, SLC35A1, UGCG, ST3GAL5*) were recently described to express endosomal receptors for HAV and other viruses (Baggen et al., 2016; Das et al., 2020; Ding et al., 2018; Drake et al., 2017; Han et al., 2018). We also identified *PTBP1* and members of the eukaryotic translation initiation factor complex (*EIF4B, EIF3C, EIF3CL*), which have known roles in IRES-mediated translation of HAV and other picornaviruses (Avanzino et al., 2017; Borman and Kean, 1997; Gosert et al., 2000; Pestova et al., 1996). The highly enriched VPS4A is an AAA-ATPase that regulates endosomal sorting complexes required for transport (ESCRT), important for budding of several enveloped viruses as well as egress of HAV (Feng et al., 2013; Votteler and Sundquist, 2013). Together, the identification of host components previously implicated in HAV infection validates the phenotypic selection in our CRISPR screen.

**Figure 1.**
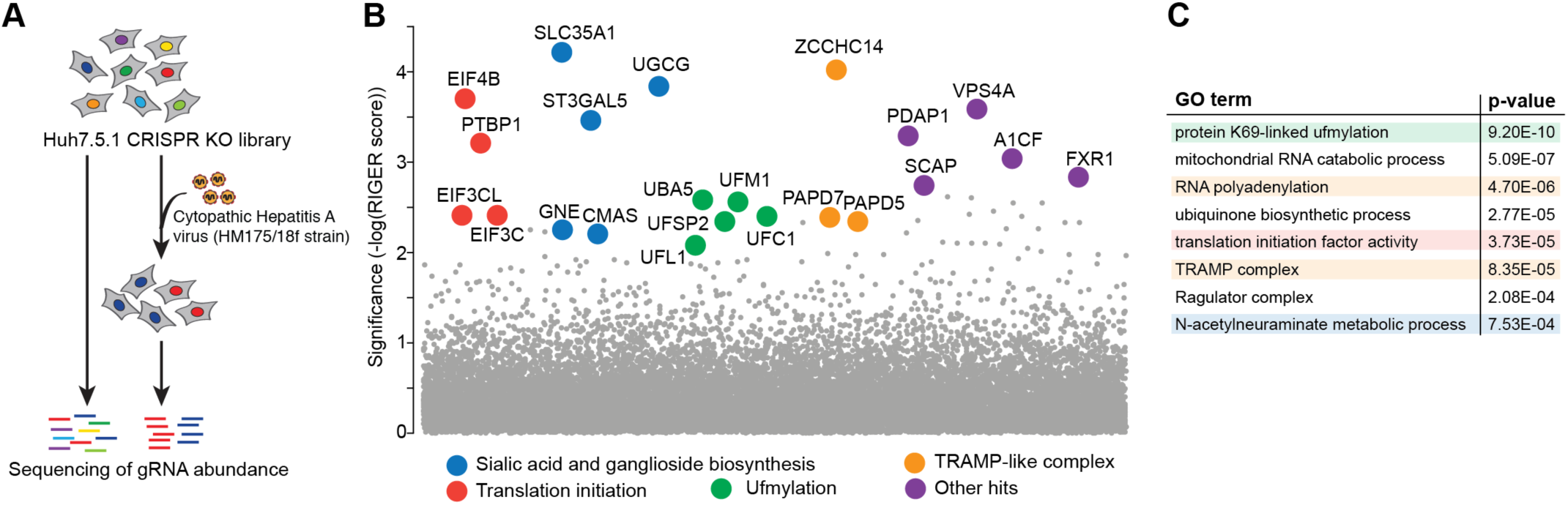
Genome-wide CRISPR KO screen identifies host cellular factors important for HAV infection. **(A)** Schematic of the genome-wide CRISPR KO screen. A library of Huh7.5.1 CRISPR-Cas9 cells was infected with cytopathic HAV HM175/18f and virus-resistant cells were harvested 12 days post-infection. gRNA abundance was measured in the original and selected cell population by next-generation sequencing. **(B)** Significance of enriched genes in CRISPR KO screen based on RIGER analysis. Hits were clustered and colored by function. **(C)** Selected terms from gene ontology (GO) analysis of genes enriched in CRISPR KO screen (RIGER p-value < 0.001).

Importantly, gene ontology (GO) analysis of enriched genes (RIGER p-value < 0.001) revealed additional pathways, which have not previously been linked to HAV infection (Figure 1C, Table S2). First, we identified all known components of the machinery for UFMylation (*UFM1, UBA5, UFL1, UFC1, UFSP2*), a ubiquitin-like protein modification (Gerakis et al., 2019; Komatsu et al., 2004) (Figures 1B and 1C). Second, the two related non-canonical poly(A) RNA polymerases, PAPD5 (also known as TENT4B or TRF4-2) and PAPD7 (also known as TENT4A or TRF4-1) were notably enriched (Figures 1B and 1C) (Warkocki et al., 2018). Finally, ZCCHC14 (Zinc Finger CCHC Domain-Containing Protein 14), the second most enriched gene, has been shown to associate with PAPD7 (Figure 1B) (Go et al., 2019). Together, these proteins form a TRAMP-like complex, generally containing an RNA polymerase and zinc finger protein, and are likely to play a critical role in HAV infection (Hamill et al., 2010; Lubas et al., 2011).

To assess the specificity of the identified cell components across different viral infections, we systematically compared our hits with CRISPR screens for host factors of human rhinovirus C15 (HRV-C15) and enterovirus D68 (EV-D68), two distantly related picornaviruses (Diep et al., 2019). Despite differences in tissue tropism and pathogenesis, HAV shares common replication strategies with other members of the *Picornaviridae* family to translate and synthesize its genomes. However, other than sialic acid biosynthesis genes among the EV-D68 hits, there were no commonalities, indicating divergent evolution of host factor dependencies (Figures S1A and S1B). Next, we compared our results to a CRISPR screen for hepatitis C virus (HCV) infection, another hepatotropic positive-sense RNA virus (and member of the *Flaviviridae* family) but found no overlap in the host factor requirements (Figure S1C) (Marceau et al., 2016).

Lastly, a comparison with results from a CRISPR screen for hepatitis B virus (HBV), a DNA virus replicating through reverse transcription, surprisingly highlighted ZCCHC14 as a shared host dependency factor between HAV and HBV (Figure S1D) (Hyrina et al., 2019). Together, these data illustrate that the host factor requirements for HAV are largely unique compared to other picornaviruses but reveal an unexpected commonality between HAV and HBV infection.

### Knockout of identified host factor genes impairs HAV replication

To validate the results of our screen, we generated isogenic KO cell lines in several genes by CRISPR-Cas9 gene editing resulting in loss of protein expression (Figures S2A-C). Upon HAV infection, we observed significant reduction of both viral replication and particle production (ranging from 10-1,000 fold) relative to levels in wild-type (WT) cells at 72 hours post-infection (hpi) (Figures 2A and S3). As PAPD5 and PAPD7 are both human orthologues of yeast Trf4p and display partial redundancy in their polyadenylation function (Warkocki et al., 2018), we generated *PAPD5/PAPD7* double knockout (DKO) cell lines (Figure S2A). We observed a significantly stronger reduction in viral replication in the DKO cells compared to the single *PAPD5* or *PAPD7* KO cells underscoring their redundant activity for HAV replication (Figure 2B). To assess viral growth phenotypes over time, we performed an infection assay using a recombinant reporter virus that expresses nanoluciferase from within its polyprotein (HAV-NLuc). We measured a ∼10^4^ fold increase in luminescence in WT cells after 72hpi but there was striking attenuation of luciferase expression in host factor KO lines, highlighting that the growth defect persists through multiple replication cycles (Figure 2C). Importantly, when we complemented *UBA5, PAPD5* and *ZCCHC14* KO cells with the respective cDNAs, we observed a significant increase in HAV replication (Figures 2D-F and S2C).

**Figure 2.**
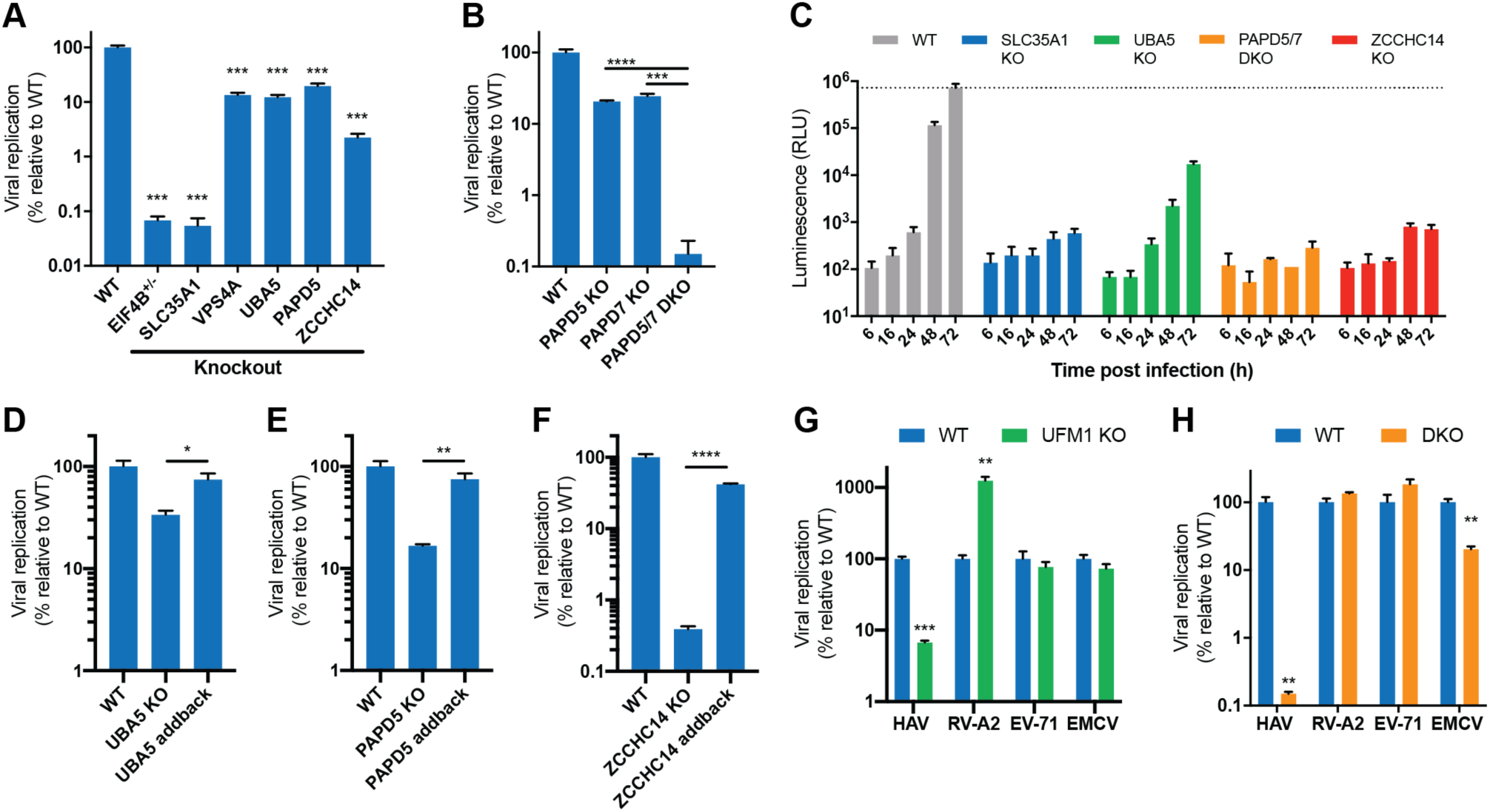
Knockout of identified host factor genes reduces HAV replication. **(A)** Quantification of HAV RNA in WT and clonal KO cell lines 3 dpi by RT-qPCR. **(B)** Quantification of HAV RNA in WT, *PAPD5* KO, *PAPD7* KO and *PAPD5/PAPD7* double KO (DKO) cells 3 dpi by RT-qPCR. **(C)** Viral growth curves using recombinant HAV expressing nanoluciferase (HAV-Nluc). WT, *SLC35A1* KO, *UBA5* KO, *PAPD5/7* DKO and *ZCCHC14* KO cells were infected, lysed at indicated timepoints and luciferase levels were measured. Dotted line indicates luminescence levels in WT cells at 72 hpi. Data are displayed as means ± s.d. (n=2 biological replicates). **(D-F)** Quantification of HAV RNA in WT, *UBA5* KO (**D**), *PAPD5* KO (**E**), or *ZCCHC14* KO (**F**) cells and KO cells complemented with respective cDNAs 3 dpi by RT-qPCR. **(G-H)** Quantification of HAV, human rhinovirus A2 (RV-A2), enterovirus 71 (EV-71) and encephalomyocarditis virus (EMCV) RNA in WT vs *UFM1* KO cells (**G**) or WT vs DKO cells (**H**) by RT-qPCR. Cells infected with HAV or RV-A2 were harvested after 48h, with EV-71 after 24h and with EMCV after 8h. For all RT-qPCR experiments, viral RNA was normalized to 18S RNA and results are displayed relative to infection in WT cells. Datasets represent means ± s.e.m. (n=3 biological replicates). P values were determined by unpaired two-sided t-tests using GraphPad Prism 8. *, p<0.05; **, p<0.01; ***, p<0.001; ****<0.0001.

It is of note that our screen did not reveal HAVCR1 (TIM1) as an essential host factor for HAV infection. To address this further, we performed a viral attachment and internalization assay in isogenic *HAVCR1* KO cells. We only observed a minor defect of internalized viral RNA at 6hpi and viral RNA levels were only reduced by ∼2 fold relative to WT cells at 30hpi (Figure S4). By contrast, in *SLC35A1* KO cells intracellular viral RNA was decreased by ∼80% at 6hpi and ∼95% at 30hpi. We conclude that *HAVCR1* is not essential for the HAV life cycle, while sialic acid biosynthesis is critical for viral entry and establishing infection. Overall, the genetic knockout studies confirm that the CRISPR screen identified genes with robust roles in the HAV life cycle.

Finally, we confirmed the finding from the comparative CRISPR screen analysis that the newly implicated UFMylation machinery and TRAMP-like complexes are distinct to HAV among different *Picornaviridae* members. HAV RNA levels were greatly reduced while enterovirus 71 (EV-71) and encephalomyocarditis virus (EMCV) RNA levels were not affected in *UFM1* KO cells (Figure 2G). Interestingly, human rhinovirus A2 (RV-A2) replication was increased suggesting that UFMylation can mediate pro- and antiviral functions in host cells, depending on the virus species. Similarly, in cells deficient in both PAPD5 and PAPD7, HAV RNA levels were severely and EMCV RNA levels moderately decreased while EV-71 and RV-A2 replication were unaffected (Figure 2H). Together, these data confirm that the genes identified by the CRISPR screen have important and distinct functions in the HAV life cycle but not for other picornaviruses.

### HAV depends on UFMylation of the ribosomal protein RPL26 for efficient translation

Struck by the unique dependency of HAV on UFMylation and no described role in viral infection, we first followed up on this cellular pathway. *UFM1* encodes the 85 amino acid long ubiquitin-fold modifier 1, which acts as a protein modification and is conserved among most eukaryotes except for yeast and other fungi (Komatsu et al., 2004). Like ubiquitination, UFM1 is covalently attached to its targets following an E1–E2–E3 reaction consisting of the UFM1-activating enzyme UBA5, the UFM1-conjugating enzyme UFC1 and the UFM1-specific ligase UFL1 (Figure 3A) (Komatsu et al., 2004; Tatsumi et al., 2010). Usually, UFM1 is present in a precursor form, which first requires cleavage of the last two C-terminal amino acids by the UFM1-specific protease UFSP2 to expose Gly at position 83 (Komatsu et al., 2004). UFSP2 also operates as the major de-UFMylation enzyme, removing UFM1 molecules from their targets (Kang et al., 2007). Our screen identified all known components of the UFMylation machinery including the additionally associated proteins DDRGK1 and ODR4 (encoded by *C1orf27*) (Figure 3A) (Chen et al., 2014; Wu et al., 2010). WT cells contain free UFM1 as well as conjugated UFM1 as detected by immunoblot (Figure 3B). To probe whether functional conjugation of UFM1 is required for HAV infection, we mutated different components of the pathway. Deletion of UBA5 led to an absence of conjugated UFM1, while UFSP2 KO resulted in hyper-conjugation, consistent with their reported activities (Figure 3B). Both conditions led to a stark reduction of viral RNA in infected cells, suggesting that the modification needs to be reversible for its proviral function (Figure 3C). Moreover, we reconstituted *UFM1* KO cells with WT UFM1, a more stable Gly83Ala UFM1 modification or ΔC3 UFM1, which lacks the last 3 C-terminal amino acids essential for conjugation (Figure 3B) (Komatsu et al., 2004). Complementation with WT UFM1 and Gly83Ala UFM1 significantly restored HAV replication (Figure 3C). By contrast, conjugation-deficient ΔC3 UFM1 did not rescue viral infection. Consistent with this, complementation of *UBA5* KO cells with WT UBA5 but not catalytically inactive enzyme also restored UFM1 conjugation and viral replication (Figures 3B and S5). Together, these data establish that functional and reversible UFM1 conjugation is critical for HAV replication.

**Figure 3.**
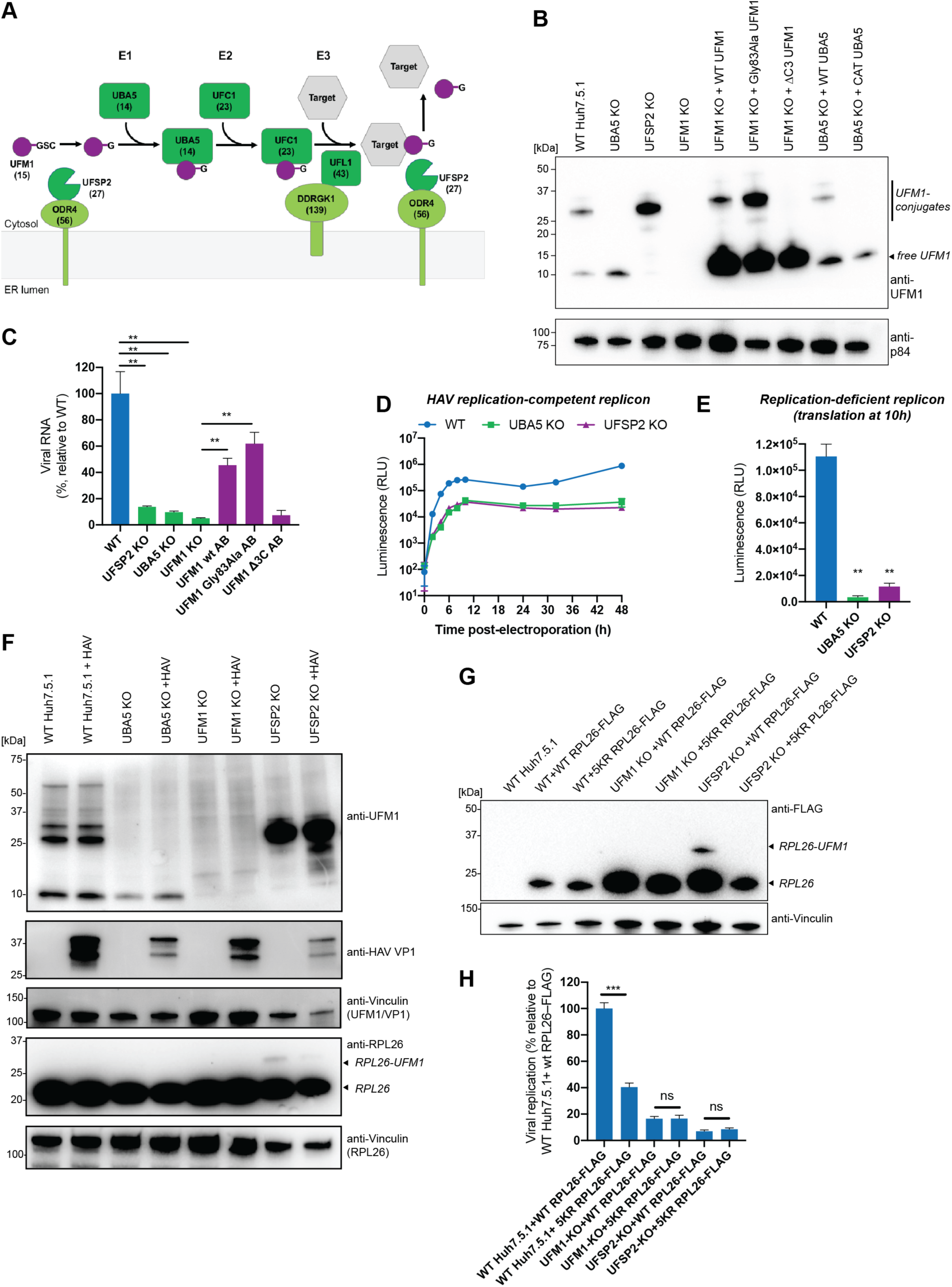
HAV depends on functional UFM1 conjugation of the ribosomal protein RPL26 for optimal translation. **(A)** Schematic of cellular UFMylation pathway. (#) indicates rank in gene enrichment analysis of CRISPR screen. **(B)** UFM1 immunoblot of WT, *UFSP2* KO, *UBA5* KO, *UFM1* KO and *UFM1* KO cells complemented with WT, Gly83Ala or ΔC3 UFM1 as well as *UBA5* KO cells complemented with WT or catalytically inactive UBA5. Upper bands represent UFM1 conjugation products and bottom band represents free UFM1. P84 was used as loading control. **(C)** Quantification of HAV RNA in WT, *UFSP2* KO, *UBA5* KO, *UFM1* KO and *UFM1* KO cells complemented with WT, Gly83Ala or ΔC3 UFM1. Viral RNA was normalized to 18S RNA and results are displayed relative to infection in WT cells. **(D)** Timecourse of luciferase activity expressed by electroporated HAV-FLuc subgenomic replicon RNA in WT, *UBA5* KO and *UFSP2* KO cells. **(E)** Luciferase activity from electroporated HAV-FLuc replication-incompetent replicon RNA in WT, *UBA5* KO and *UFSP2* KO cells at 10h post-electroporation. **(F)** Immunoblots of uninfected and HAV infected WT, *UBA5* KO, *UFM1* KO and *UFSP2* KO cells. Membranes were stained with anti-UFM1, anti-HAV VP1 or anti-RPL26 antibodies. Anti-Vinculin served as loading control. **(G)** FLAG immunoblot of WT, *UFM1* KO and *UFSP2* KO cells transduced with lentivirus expressing FLAG-tagged WT RPL26 or 5KR RPL26. Vinculin served as loading control. **(H)** Quantification of HAV RNA in WT, *UFM1* KO and *UFSP2* KO cells expressing WT RPL26 or 5KR RPL26. Viral RNA was normalized to 18S RNA and results are displayed relative to infection in WT cells expressing WT RPL26. For all RT-qPCR experiments, datasets represent means ± s.e.m. (n=3 biological replicates). For luciferase experiments, datasets represent means ± s.d. (n=2 biological replicates) for each timepoint. P values were determined by unpaired two-sided t-tests using GraphPad Prism 8. ns, non-significant; *, p<0.05; **, p<0.01; ***, p<0.001; ****<0.0001.

Next, in order to assess whether the defect occurs during early viral translation or later during the genome replication step, we used HAV replicon RNA expressing firefly luciferase (Fluc). We detected a marked defect in *UBA5* and *UFSP2* KO cells within 10h post-electroporation (Figure 3D). To specifically measure the effect on translation of incoming viral RNA, we used a replication-incompetent replicon (containing a GDD-->GAA mutation in the viral polymerase) and observed strongly reduced luciferase expression in both KO cells (Figures 3E and S6A). We excluded that observed phenotypes were due to differences in cell viability or proliferation post-electroporation (Figure S6B). We therefore conclude that UFMylation is required for optimal translation of HAV RNA.

Recently, the ribosomal protein RPL26 was identified as a major target for UFMylation in human cells (Walczak et al., 2019; Wang et al., 2019). Thus, we tested whether UFMylated RPL26 is important for efficient HAV translation. Consistent with previous findings, we detected modified RPL26 at the molecular weight corresponding to UFM1 conjugates (Figure 3F). Interestingly, only a small fraction of RPL26 is UFMylated and the conjugation appears transient as it can only be visualized in *UFSP2* KO cells, which lack de-UFMylation activity. It was further shown that conjugation of UFM1 occurs at C-terminal lysines of RPL26 (Walczak et al., 2019; Wang et al., 2019). We generated cell lines with ectopic WT RPL26 or cDNA containing five lysine to arginine mutations (“5KR RPL26”) and confirmed that ectopic 5KR RPL26 was not UFMylated (Figures 3G and S7A). HAV infection of WT Huh7.5.1 cells expressing 5KR RPL26 was reduced compared to cells containing ectopic WT RPL26, which largely phenocopied the *UFM1* KO effect (Figure 3H). By contrast, there was no dominant negative effect of 5KR RPL26 in transduced *UFM1* or *UFSP2* KO cells, where HAV replication was already fully reduced compared to WT Huh7.5.1 containing WT RPL26 (Figure 3H). Similarly, we measured reduced HAV RNA levels in previously characterized U2OS cells expressing 5KR RPL26 and a CRISPRi system to knockdown endogenous WT RPL26 (Walczak et al., 2019) (Figure S7B). Lastly, 5KR RPL26 cells exhibited increased RV-A2 infection (Figure S7C), thus mirroring the effect of *UFM1* deletion (Figure 2G). Together, the results indicate that UFM1 and RPL26 are functionally linked and support a model where HAV requires active UFMylation of RPL26 for translation.

### TRAMP-like complexes support HAV translation independent of polyadenylation of HAV tails

Intrigued by the strong phenotypes on HAV replication and a critical role in the life cycles of both HAV and HBV, we performed mechanistic studies on the TRAMP-like complex components PAPD5, PAPD7 and ZCCHC14. The two non-canonical poly(A) RNA polymerases PAPD5 and PAPD7 can transfer poly or oligo(A) to a variety of RNA substrates (e.g. mRNAs, miRNAs, snoRNAs and rRNAs) in a template-independent manner (Berndt et al., 2012; Boele et al., 2014; Rammelt et al., 2011; Shcherbik et al., 2010; Shin et al., 2017). Their cellular activities include stabilization of mRNAs for translation as well as specifying different RNA species for turnover by the exosome complex (Lubas et al., 2011; Vaňáčová et al., 2005; Warkocki et al., 2018). Recently, ZCCHC14 has been shown to interact with PAPD5 and PAPD7 to support HBV infection (Hyrina et al., 2019) and consistent with that we observed strong reduction of HBV RNA levels in *ZCCHC14* KO cells (Figure S8A).

To determine the function of the TRAMP-like complex during HAV infection, we first measured the effects of *PAPD5/7* DKO or *ZCCHC14* KO on viral translation and replication using viral replicon RNA. We observed a notable defect in luciferase expression immediately post-electroporation, suggesting inefficient translation of viral RNA (Figure 4A). While WT cells supported replicon replication at later timepoints, luminescence decayed in the KO cells. We additionally probed viral translation directly using the replication-incompetent HAV replicon and observed drastically reduced luciferase expression from incoming RNA immediately post-electroporation in DKO and *ZCCHC14* KO cells (Figures 4B). By contrast, translation of electroporated firefly mRNA was not affected, excluding a general translation defect in KO cells (Figure S8B). As translation from viral RNA is directly linked to RNA stability, we measured HAV RNA levels upon infection over time. However, we did not detect differences in viral RNA levels between WT, DKO or *ZCCHC14* KO cells early during infection (Figure 4C). We therefore conclude that deletion of TRAMP-complex components affects HAV translation but not HAV RNA stability.

**Figure 4.**
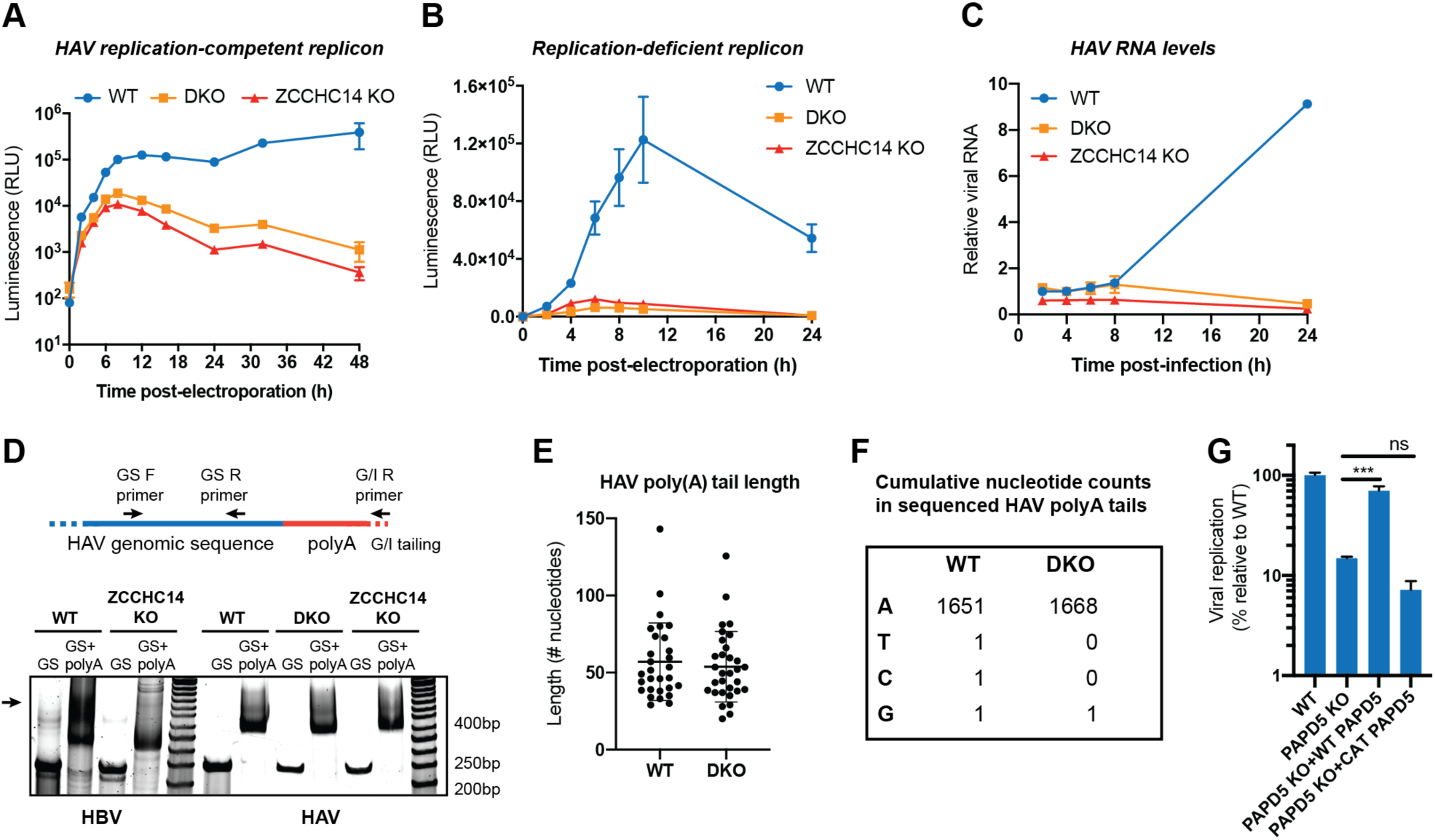
Catalytic activity of TRAMP-like complexes is important for HAV translation, independent of regulating viral poly(A) tail length or RNA stability. **(A)** Timecourse of luciferase activity expressed by electroporated HAV-FLuc subgenomic replicon RNA in WT, DKO and *ZCCHC14* KO cells. Datasets represent means ± s.d. (n=2 biological replicates) for each timepoint. **(B)** Timecourse of luciferase activity expressed by electroporated HAV-FLuc replication-incompetent replicon RNA in WT, DKO cells and *ZCCHC14* KO. Datasets represent means ± s.d. (n=2 biological replicates) for each timepoint. **(C)** Quantification of HAV RNA in WT, DKO and *ZCCHC14* KO cells by RT-qPCR. Cells were infected on ice, incubated at 37C for 1h, washed 3 times and harvested at indicated times. Viral RNA was normalized to 18S RNA. Datasets represent means ± s.e.m. (n=3 biological replicates). **(D)** Poly(A) tail length assay. Total RNA from infected WT or KO cells was G/I tailed at 3’ends, reverse transcribed and PCR amplified by either genome-specific (GS) forward and GS reverse primers, or GS forward and G/I reverse primer. PCR products were analyzed by polyacrylamide gel electrophoresis. For HBV, total RNA was collected from infected WT or *ZCCHC14* KO HepG2-NTCP cells 14dpi. For HAV, total RNA was harvested from infected WT, DKO or *ZCCHC14* KO Huh7.5.1 cells 3dpi. Arrow indicates a longer HBV poly(A) tail in WT cells. **(E)** Length distribution of sequenced HAV poly(A) tails. An RNA linker was ligated to 3’ends of rRNA-depleted RNA from infected WT or DKO cells followed by reverse transcription and PCR amplification of the HAV poly(A) tail region. PCR products were TOPO cloned and Sanger sequenced. **(F)** Analysis of nucleotide composition of sequenced HAV poly(A) tails from (E). **(G)** Quantification of HAV RNA in WT, *PAPD5* KO and *PAPD5* KO cells complemented with either WT or catalytically inactive (CAT) PAPD5 3 dpi by RT-qPCR. Viral RNA was normalized to 18S RNA and results are displayed relative to infection in WT cells. Datasets represent means ± s.e.m. (n=3 biological replicates).

For HBV, mechanistic studies revealed that HBV surface antigen expression requires RNA tailing by TRAMP-like complexes (Hyrina et al., 2019). As HAV has a polyadenylated genome and longer poly(A) tails can promote translation (Bergamini et al., 2000), we assessed the length distribution of HAV poly(A) tails from infected WT and KO cells by polyacrylamide gel electrophoresis but we did not observe a notable difference (Figure 4D). By contrast, HBV tails were reduced in *ZCCHC14* KO cells, confirming the expected defect (Figure 4D). To corroborate that HAV tail length is not affected, we performed TOPO cloning followed by Sanger sequencing as well as a higher resolution electrophoretic shift assay using TapeStation, confirming that HAV tails are not shortened in KO cells (Figures 4E and S8C).

To explore a different function of PAPD5 and PAPD7 for HAV infection, we looked at the composition of viral poly(A) tails. Recently, it was shown that PAPD5 and PAPD7 can introduce guanosines at low frequencies into poly(A) tails resulting in increased stability against CCR4-NOT dependent deadenylation (Lim et al., 2018). However, we did not observe an elevated frequency of non-adenosine nucleotides in WT vs DKO cells within HAV poly(A) tails (Figure 4F). Rather, almost all analyzed tails exclusively consisted of adenosines in both conditions. Together, the data suggest that TRAMP-like complexes do not support viral translation by regulating the length or composition of HAV poly(A) tails.

As the described cellular functions of PAPD5 and PAPD7 depend on their catalytic activity, we complemented KO or DKO cells with WT or catalytically inactive (CAT) (containing two aspartate to alanine substitutions in the active site) cDNAs (Figure S2C) (Rammelt et al., 2011). Importantly, infection of the complemented cell lines established that the catalytic activities of PAPD5 and PAPD7 are required to support viral replication (Figures 4G and S8D). We speculate that the TRAMP-like complex either modulates HAV RNA in a (so far) unrecognized manner or that expression of a cellular protein, which is critical for HAV translation, depends on polyadenylation of its transcript. In conclusion, we find that the catalytic activity is required to support viral translation, independent of directly modulating viral poly(A) tails or RNA stability.

### Pharmacological Inhibition of PAPD5 and PAPD7 exhibits an antiviral effect against HAV infection

As the enzymatic function of PAPD5 and PAPD7 are required for HAV infection, we explored whether pharmacological inhibition has an antiviral effect. We used RG7834 (Figure 5A), a small molecule compound belonging to the chemical class of the dihydroquinolizinones. Interestingly, this compound was first identified as a potent inhibitor for HBV surface antigen secretion (Mueller et al., 2018), and PAPD5 and PAPD7 were subsequently shown to be its cellular targets (Mueller et al., 2019).

**Figure 5.**
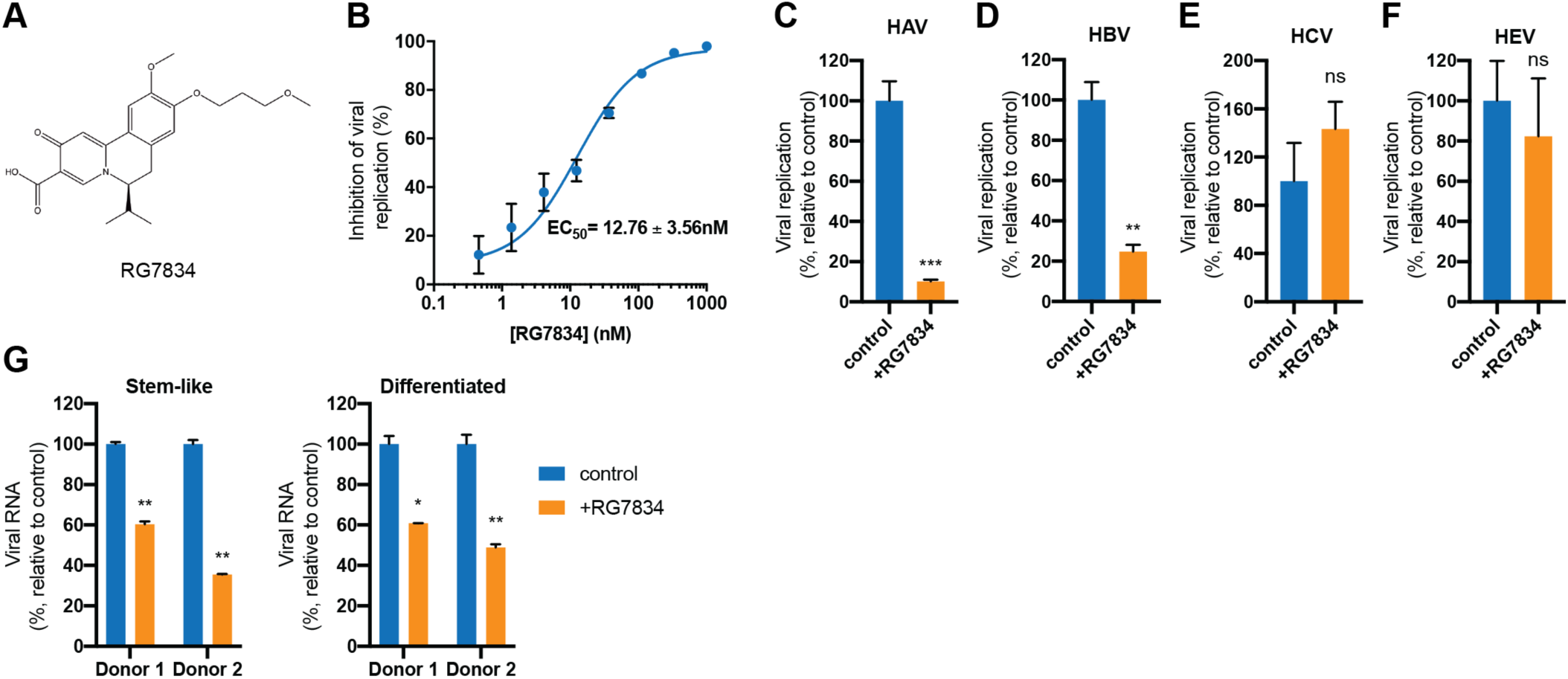
Small molecule inhibition of PAPD5 and PAPD7 reduces HAV infection in human hepatocyte cells and human liver organoids. **(A)** Chemical structure of RG7834. **(B)** Dose-response curve of HAV RNA levels in Huh7.5.1 cells treated with different RG7834 concentrations as measured by RT-qPCR 3 days post-infection/treatment. Non-linear curve was fitted with least squares regression using GraphPad Prism 8 and EC_50_ was determined. **(C)** Quantification of HAV RNA in control and RG7834 (100 nM) treated Huh7.5.1 cells 3 dpi by RT-qPCR. **(D)** Quantification of HBV total RNA in control and RG7834 (100 nM) treated HepG2-NTCP cells 15 dpi by RT-qPCR. Drug was replenished every 3-4 days. **(E)** Quantification of HCV RNA in control and RG7834 (100 nM) treated Huh7.5.1 cells 3 dpi by RT-qPCR. **(F)** Quantification of HEV RNA in control and RG7834 (100 nM) treated Huh7.5.1 cells by RT-qPCR 6 days post-transfection of viral RNA. **(G)** Quantification of HAV RNA in control and RG7834 (100 nM) treated human liver organoids 7 dpi by RT-qPCR. Organoids were either stem-like or differentiated to adult hepatocytes prior to infection. For all RT-qPCR experiments, viral RNA was normalized to 18S RNA and results are displayed relative to infection in control (DMSO) condition. Datasets in (**B-F**) represent means ± s.e.m. (n=3 biological replicates). For organoid experiments (**G**) donor 1 and 2 represent 2 independent biological replicates and qPCR was performed with 2 technical replicates. P values were determined by unpaired two-sided t-tests using GraphPad Prism 8. ns, non-significant; *, p<0.05; **, p<0.01; ***, p<0.001.

To test whether RG7834 also exhibits activity against HAV replication, we performed a dose-response curve and calculated a 50% effective concentration (EC_50_) of 12.76 nM for inhibition of HAV replication (Figure 5B). This is far below the measured 50% cytotoxic concentration (CC_50_) of >10 µM (Figure S9A). Drug treatment of infected *PAPD5* or *PAPD7* KO cells suggested that RG7834 has a stronger inhibitory effect on the PAPD5 activity as reduction of viral RNA is more substantial in the *PAPD7* KO+RG7834 condition (Figure S9B). Intrigued by the fact that RG7834 inhibits both HAV and HBV, we also measured the effects on two additional liver-specific viruses, HCV and hepatitis E virus (HEV). However, RG7834 only showed significant reduction of HAV and HBV RNA levels (Figure 5C-F). This is consistent with the comparative CRISPR screen analysis for HCV as well as unchanged viral replication levels in DKO cells for both HCV and HEV (Figure S9C).

Finally, we performed infection assays in human liver organoid cultures (Broutier et al., 2016). We observed significant dose-dependent reduction of HAV RNA in both stem cell-like and differentiated organoids under RG7834 treatment (Figures 5G and S9D). Together, our results demonstrate that pharmacological inhibition of PAPD5/7 is a promising strategy for host-directed therapy of HAV infection.

## Discussion

This study provides a comprehensive assessment of the host factors that are critical for HAV infection. The identified genes comprise several cellular pathways and complexes, such as sialic acid biosynthesis, eukaryotic translation initiation, UFMylation, TRAMP-like complexes and others. We corroborated that the putative HAV receptor, HAVCR1, is not essential for infection. Rather, we found multiple genes linked to sialic acid and ganglioside biosynthesis, which are important for efficient HAV uptake (Das et al., 2020). The screen also identified eukaryotic translation initiation factor components and PTBP1, which were previously linked to HAV translation (Avanzino et al., 2017; Borman and Kean, 1997; Gosert et al., 2000; Pestova et al., 1996).

Importantly, we revealed two cellular pathways, UFMylation and TRAMP-like complexes, which had not been implicated in HAV life cycle before. UFMylation is a ubiquitin-like, post-translation modification. We demonstrated that HAV specifically depends on UFMylation of the ribosomal protein RPL26 for efficient translation. The cellular function of this protein modification is still an emerging field of research but it has been linked to the regulation of the ER stress response and ribosome-associated quality control (Walczak et al., 2019; Wang et al., 2019). Specific ribosomal or ribosome-associated proteins have been shown to be required for efficient translation of other viral genomes (Landry et al., 2009; Majzoub et al., 2014). RPL26 is located near the ribosome exit tunnel and its modification could facilitate dynamic and reversible conformation changes to improve translation of specific amino acid sequences or resolve certain secondary structure in the RNA as suggested by the requirement for active UFMylation and de-UFMylation during HAV infection. Additional studies, such as biochemical analysis and ribosome profiling of the different ribosomal subpopulations, are needed to gain a better understanding of its role during translation of viral and cellular mRNAs. The contrasting phenotypes observed during HAV and RV-A2 infection in *UFM1* KO cells may also be utilized as tools for future studies of the molecular effects of UFMylation.

Furthermore, we identified the TRAMP-like complex components ZCCHC14, PAPD5 and PAPD7 and demonstrated that they are required for HAV translation. PAPD5 and PAPD7 have been described to possess polyadenylation activity as well as the ability to generate mixed tails (Lim et al., 2018; Warkocki et al., 2018). Despite showing that their catalytic activity is required to support HAV infection, we did not observe any abnormalities in HAV poly(A) tail length, composition or viral RNA stability in TRAMP-like complex deficient cells. However, this is consistent with previous observations that picornavirus polyadenylation is template-dependent, for example by the presence of a homotypic poly(U) sequence in the picornavirus negative-sense RNA (Spector and Baltimore, 1975). Moreover, studies using poliovirus, another picornavirus, suggested that the picornaviral RNA-dependent RNA polymerase uses reiterative transcription as it replicates the poly(A) tails of viral RNA (Kempf et al., 2013; Steil et al., 2010). These findings support that PAPD5 and PAPD7 are not necessarily required for HAV poly(A) tail generation or maintenance. Thus, the TRAMP-like complex may fulfill other functions, for example by directly regulating viral translation. In studies using *C. elegans*, deletion of the worm orthologs of PAPD5 (gld-4) and ZCCHC14 (gls-1) led to reduced polysome formation but did not affect global poly(A) tail lengths, providing evidence of polyadenylation-independent functions of TRAMP-like complexes (Nousch et al., 2014). Alternatively, PAPD5 and PAPD7 could regulate the expression of another host factor critical for HAV translation through RNA tailing.

Interestingly, other tested picornaviruses were not dependent on UFMylation or the TRAMP-like complex for efficient translation. Therefore, our data suggests that HAV translation differs even more substantially from other picornaviruses than previously recognized. While it is known that HAV is distinct in its dependency on the cellular eukaryotic translation initiation factor complex (Borman and Kean, 1997), we additionally revealed an unique requirement of UFMylation and the TRAMP-like complex for HAV.

Surprisingly though, PAPD5, PAPD7 and ZCCHC14 were identified as critical factors for infection with HBV, a hepatotropic DNA virus with a vastly different life cycle, using chemical and genetic screens (Hyrina et al., 2019; Mueller et al., 2019). Mechanistically, the TRAMP-like complex plays an important role in HBV surface antigen expression by poly(A) tail extension of HBV mRNAs (Hyrina et al., 2019). This highlights a remarkable convergence of two unrelated hepatitis viruses to co-opt the same cellular complex in different ways.

Finally, based on these molecular insights, we provided proof-of-concept that pharmacological inhibition of PAPD5 and PAPD7 decreases infection levels of HAV and HBV. Thus, this study exemplifies that host factor screens are a powerful strategy to gain a comprehensive understanding of the most critical cellular pathways for viral infections, and to illuminate new drug targets, which can be shared across different viral families.

---

**Figure S1.**
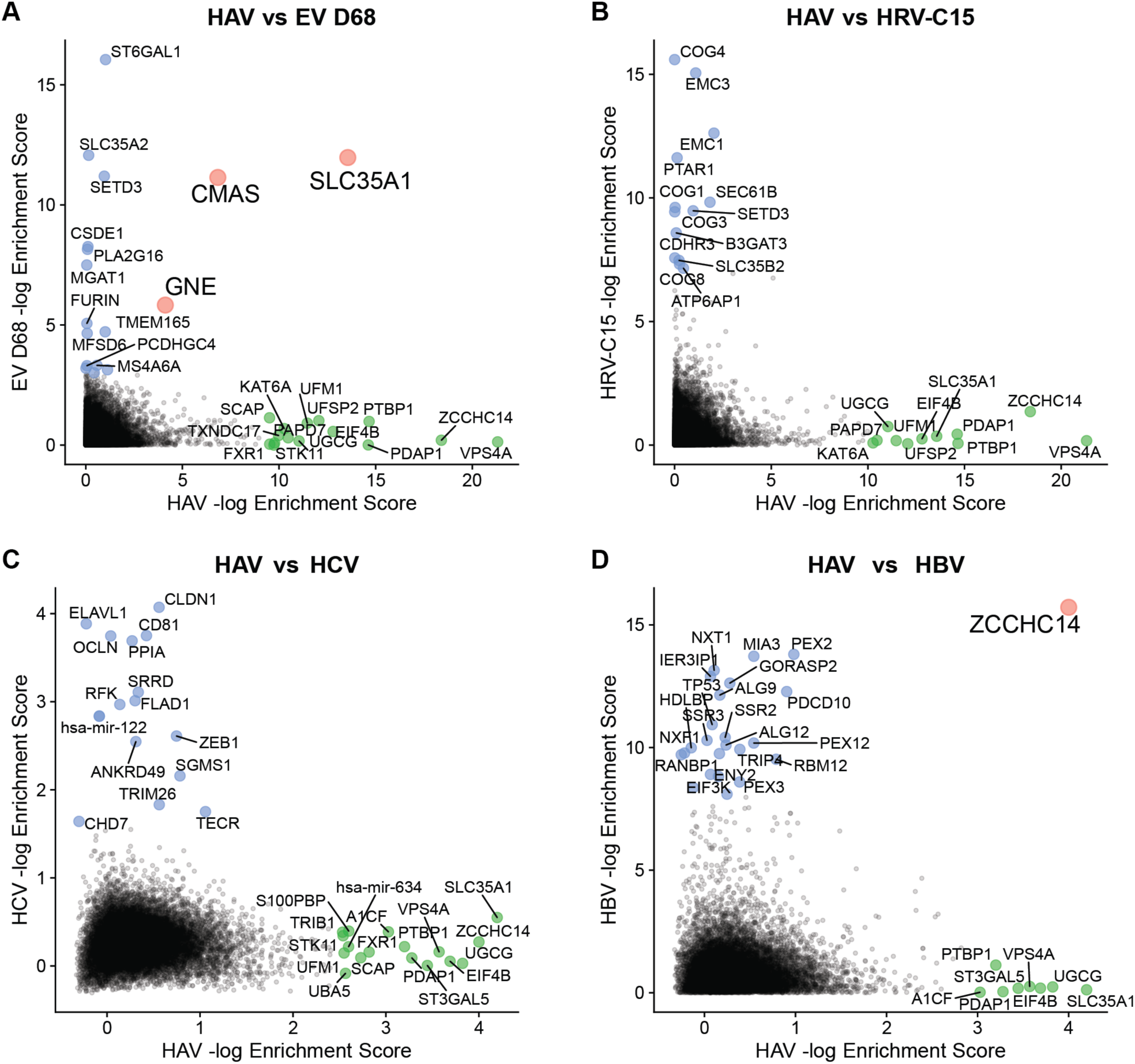
Comparative CRISPR screen analysis reveals that HAV host factors are distinct from other picornaviruses and HCV but highlights that ZCCHC14 is a shared hit between HAV and HBV, Related to Figure 1. Comparison of HAV CRISPR screen gene enrichment scores with hits from **(A)** EV-D68 (Diep et al., 2019), **(B)** HRV-C15 (Diep et al., 2019), **(C)** HCV (Marceau et al., 2016), or **(D)** HBV CRISPR screens (Hyrina et al., 2019). Overlapping hits from both screens are highlighted in red.

**Figure S2.**
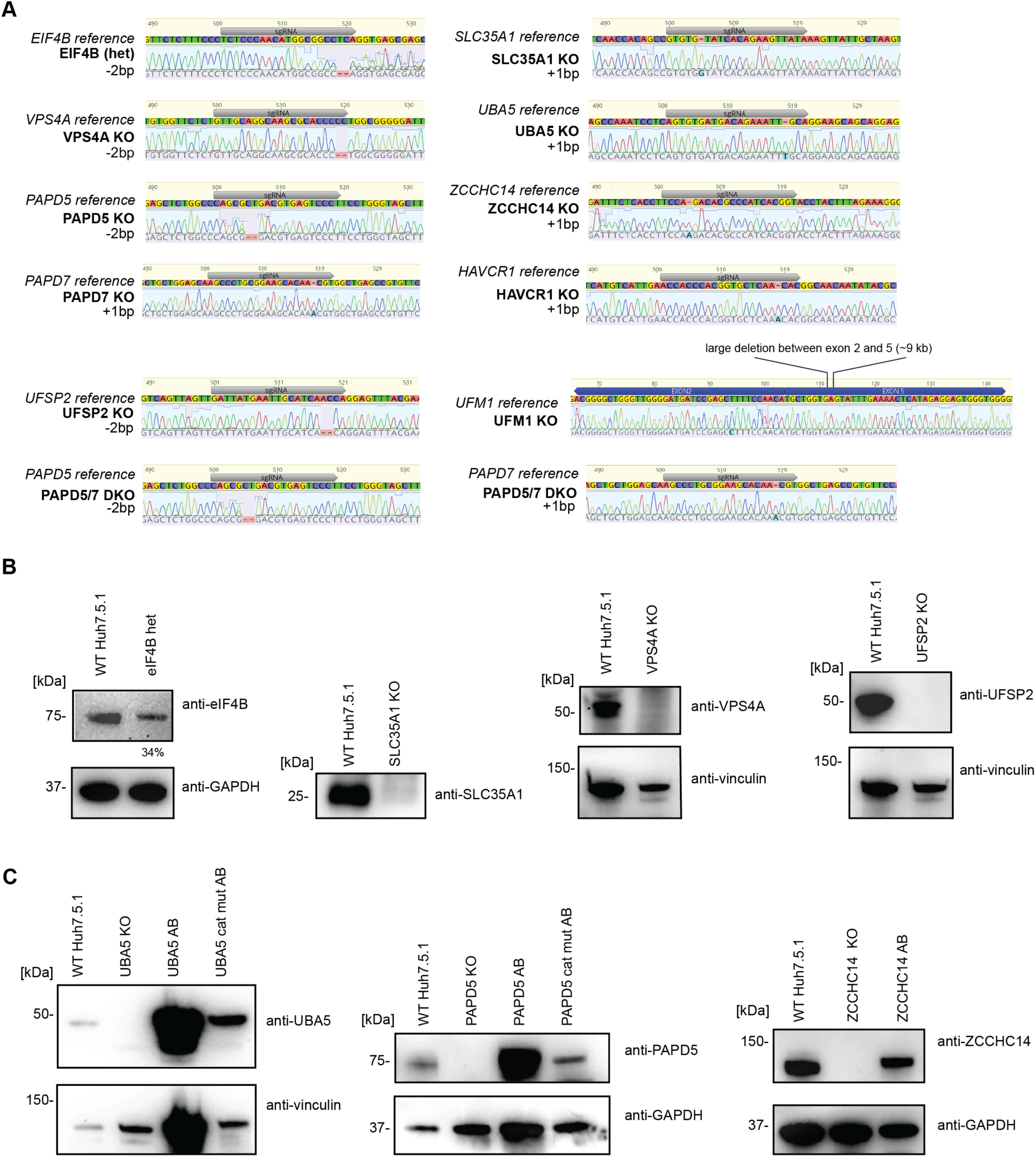
Genotyping and immunoblotting of clonal Huh7.5.1 KO cell lines, Related to Figure 2. **(A)** Sanger sequencing of gRNA targeted loci of clonal KO cell lines. Traces were aligned to corresponding reference sequences and indel mutations determined. Reads of gRNA targeted PAPD5 and PAPD7 loci in clonal Huh7.5.1 DKO cells are displayed. UFM1 gene was simultaneously targeted by two gRNAs cutting in exon 2 and exon 5 resulting in a large genomic deletion. **(B)** Immunoblots for clonal *eIF4B, SLC35A1, VPS4A* and *UFSP2* KO cell lines. For eIF4B, a heterozygous clone harboring one allele with a −2bp deletion and one WT allele, quantification of the band (in %) is normalized to GAPDH signal and relative to WT indicating decrease of expression in heterozygous clone. For SLC35A1, WT and KO cells were lysed in RIPA buffer and lysates were immunoprecipitated using anti-SLC35A1 antibody and protein G beads prior to immunoblot. GAPDH and Vinculin were used as loading controls. **(C)** Immunoblots of WT, *UBA5* KO, *PAPD5* KO or *ZCCHC14* KO cells as well as KO cells complemented with respective cDNAs. GAPDH and Vinculin were used as loading controls.

**Figure S3.**
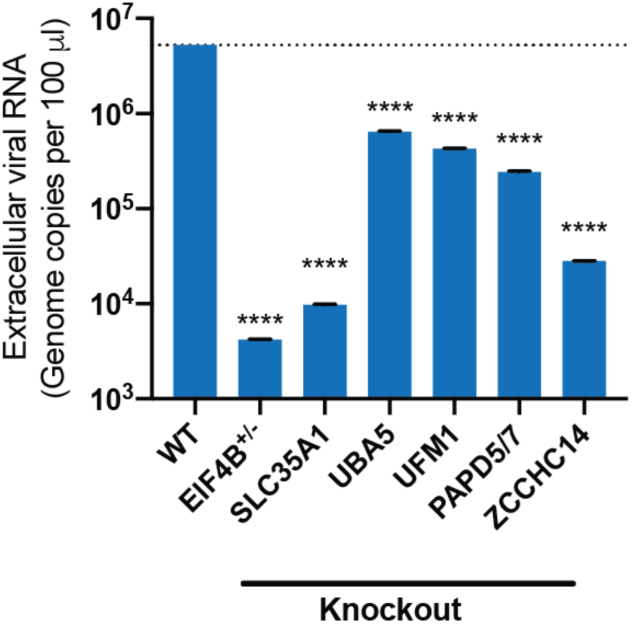
Host factor knockout decreases extracellular viral RNA, Related to Figure 2. Quantification of extracellular viral RNA in supernatant of cells 3 dpi using HM175/18f HAV. RNA was extracted from supernatant, measured by RT-qPCR and compared to a standard curve of in vitro transcribed viral RNA to determine genome copies. Dataset represents means ± s.e.m. (n=2 biological replicates). P values were determined by one-way Anova with uncorrected Fisher’s Least Significant Difference test using GraphPad Prism 8. *, p<0.05; **, p<0.01; ***, p<0.001; ****<0.0001.

**Figure S4.**
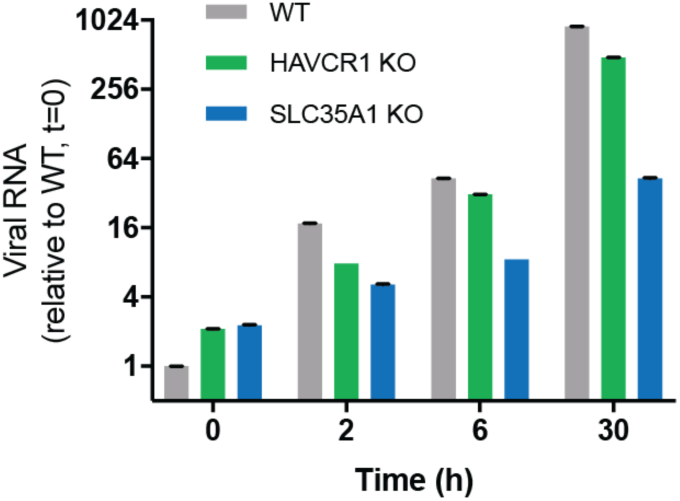
Deletion of sialic acid biosynthesis severely impairs viral entry, while *HAVCR1* KO is not critical for viral uptake, Related to Figure 2. Quantification of HAV RNA in WT, *HAVCR1* KO and *SLC35A1* KO cells by RT-qPCR after cells were infected on ice, incubated at 37C, and washed before harvest at indicated times. Viral RNA was normalized to 18S RNA and results are displayed relative to infection in WT cells. Datasets represent means ± s.e.m. (n=3 biological replicates).

**Figure S5.**
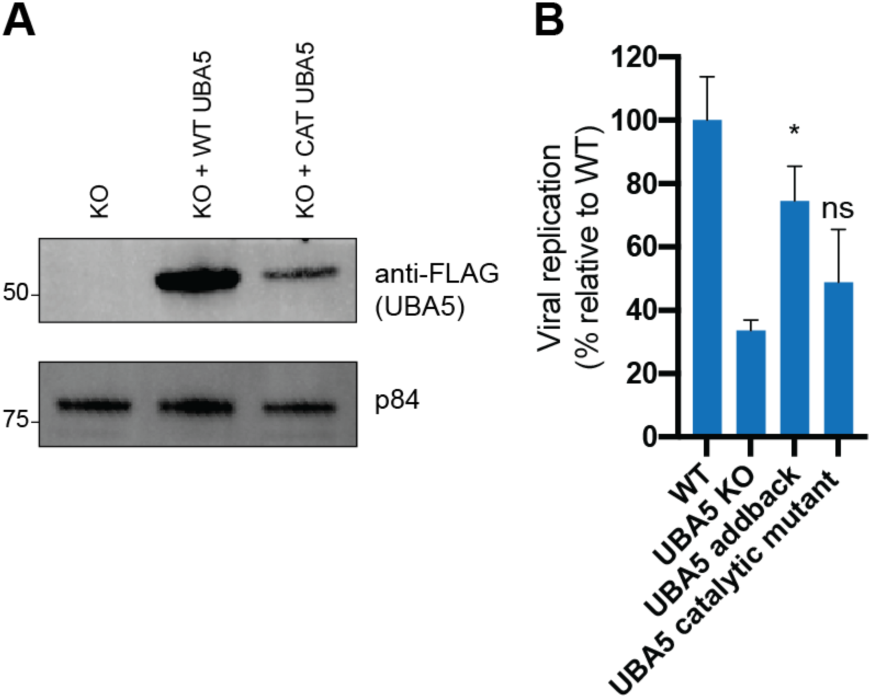
HAV depends on catalytically active UBA5, Related to Figure 3. **(A)** FLAG immunoblot of lysates from *UBA5* KO cells transduced with FLAG-tagged WT or catalytically inactive (CAT) UBA5. P84 was used as loading control. **(B)** Quantification of HAV RNA in WT, *UBA5* KO and *UBA5* KO complemented with WT or CAT UBA5 3 dpi by RT-qPCR. Viral RNA was normalized to 18S RNA and results are displayed relative to infection in WT cells. Datasets represent means ± s.e.m. (n=3 biological replicates). P values were determined by unpaired two-sided t-tests using GraphPad Prism 8. ns, non-significant; *, p<0.05.

**Figure S6.**
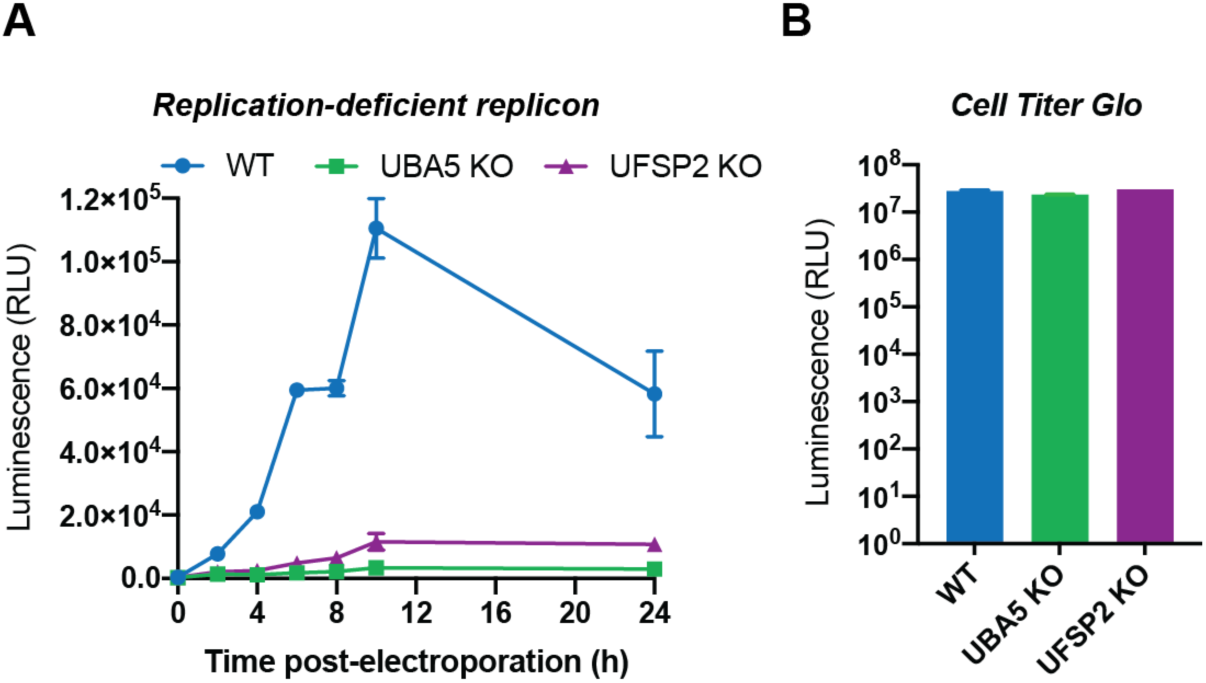
HAV translation is impaired in UFMylation deficient cells, Related to Figure 3. **(A)** Timecourse of luciferase activity expressed by electroporated HAV-FLuc replication-incompetent replicon RNA in WT, *UBA5* KO and *UFSP2* KO cells. Datasets represent means ± s.d. (n=2 biological replicates) for each timepoint. **(B)** Cell Titer Glo assay to measure cell viability of electroporated WT, *UBA5* KO and *UFSP2* KO cells 24h post-electroporation. Datasets represent means ± s.d. (n=2 biological replicates).

**Figure S7.**
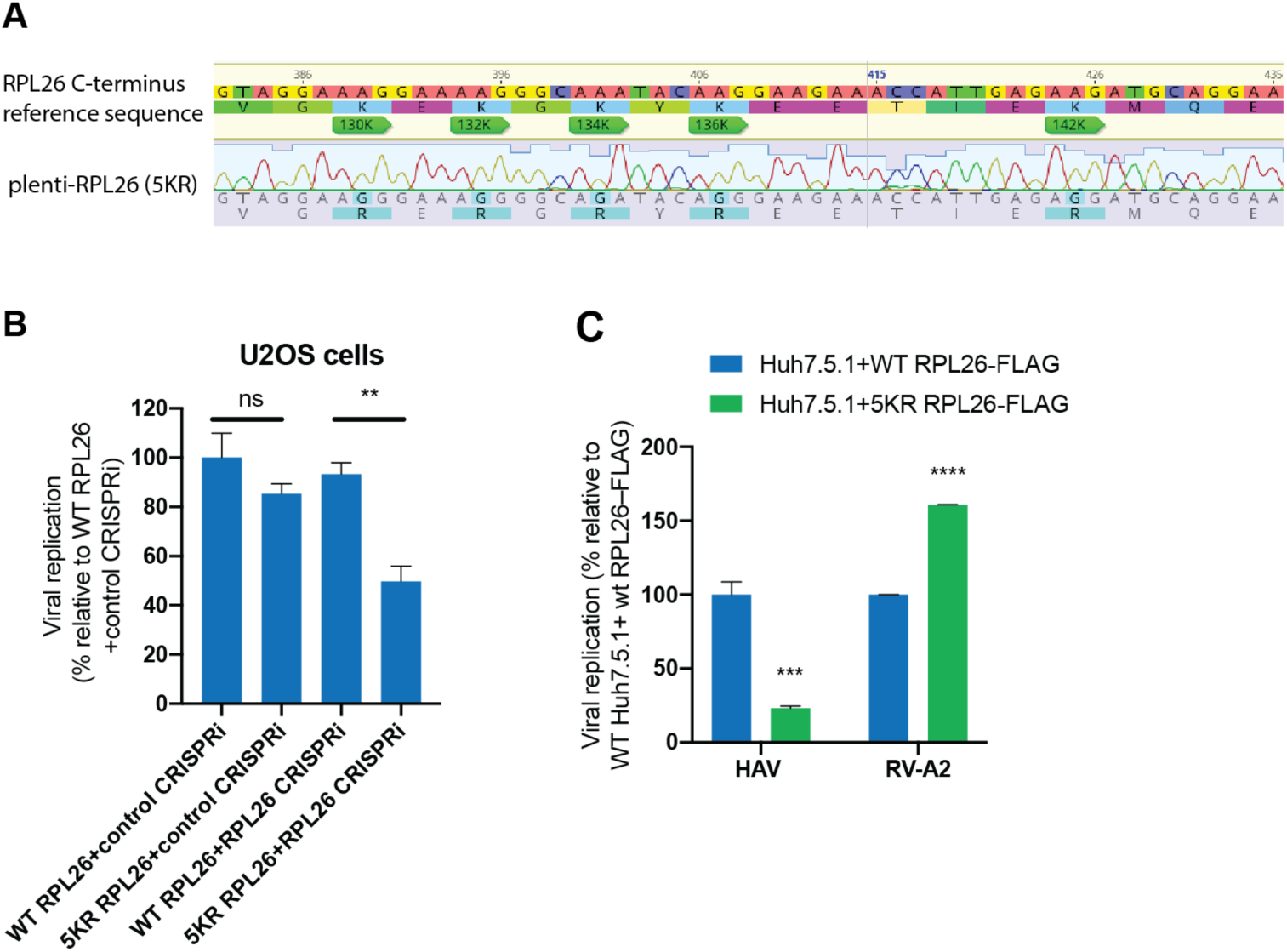
UFM1 conjugation to C-terminal lysines of RPL26 is required for optimal HAV infection, Related to Figure 3. **(A)** Reference and Sanger sequence of lentiviral construct expressing RPL26 with five lysine to arginine point mutations at C-terminus (5KR). **(B)** Quantification of HAV RNA in U2OS dCas9-KRAB cells expressing WT RPL26 or 5KR RPL26 as well as a control (non-targeting) or RPL26 targeting gRNA. Viral RNA was normalized to 18S RNA and results are displayed relative to infection in cells expressing WT RPL26 and the control gRNA. **(C)** Quantification of HAV and RV-A2 RNA in WT cells transduced with lentivirus expressing FLAG-tagged WT RPL26 or 5KR RPL26. Viral RNA was normalized to 18S RNA and results are displayed relative to infection in WT cells expressing WT RPL26. Dataset represents means ± s.e.m. (n=3 biological replicates). P values were determined by unpaired two-sided t-tests using GraphPad Prism 8. ns, non-significant; *, p<0.05; **, p<0.01; ***, p<0.001; ****<0.0001.

**Figure S8.**
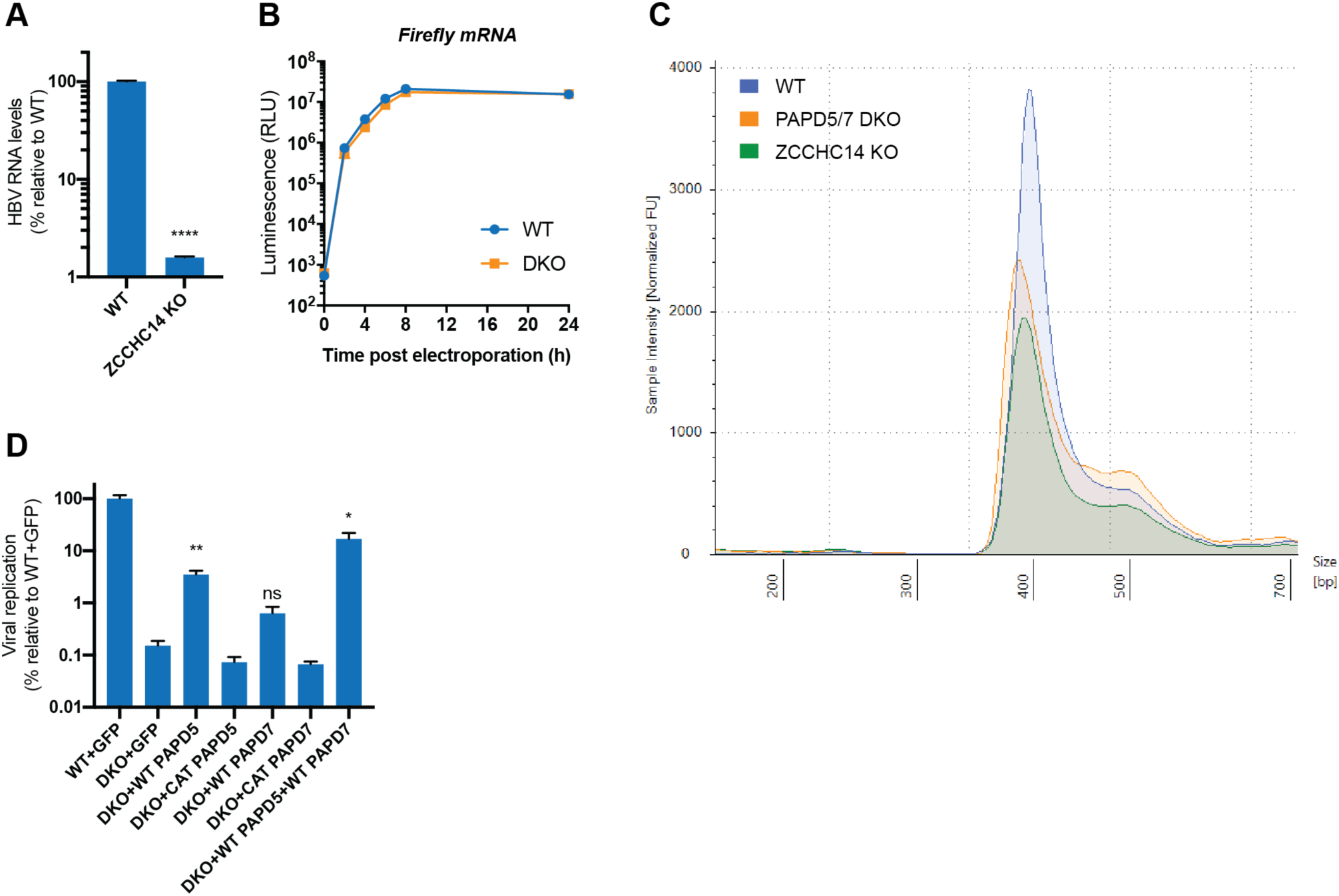
Catalytic activity of TRAMP-like complexes is important for HAV translation, independent of regulating viral poly(A) tail length or RNA stability, Related to Figure 4. **(A)** Quantification of HBV RNA in WT and *ZCCHC14* KO HepG2-NTCP cells 14dpi. Viral RNA was normalized to 18S RNA and results are displayed relative to infection in WT cells. **(B)** Timecourse of luciferase activity expressed by electroporated firefly mRNA in WT and DKO cells. Datasets represent the means ± s.d. (n=2 biological replicates). **(C)** Electrophoretic analysis of poly(A) tail region of HAV from infected WT, DKO and *ZCCHC14* KO cells at 72hpi. Total RNA was G/I tailed at 3’ends, reverse transcribed and PCR amplified by forward primer ∼300bp upstream of HAV poly(A) start site and G/I specific reverse primer. PCR products were analyzed using TapeStation. **(D)** Quantification of HAV RNA in DKO cells transfected with GFP cDNA, PAPD5 WT or CAT cDNA, PAPD7 WT or CAT cDNA, or co-transfected with both PAPD5 and PAPD7 WT cDNAs. Viral RNA was normalized to 18S RNA and results are displayed relative to infection in WT cells transfected with GFP cDNA. For RT-qPCR, datasets represent means ± s.e.m. (n=3 biological replicates). P values were determined by unpaired two-sided t-tests using GraphPad Prism 8. ns, non-significant; *, p<0.05; **, p<0.01; ****, p<0.0001.

**Figure S9.**
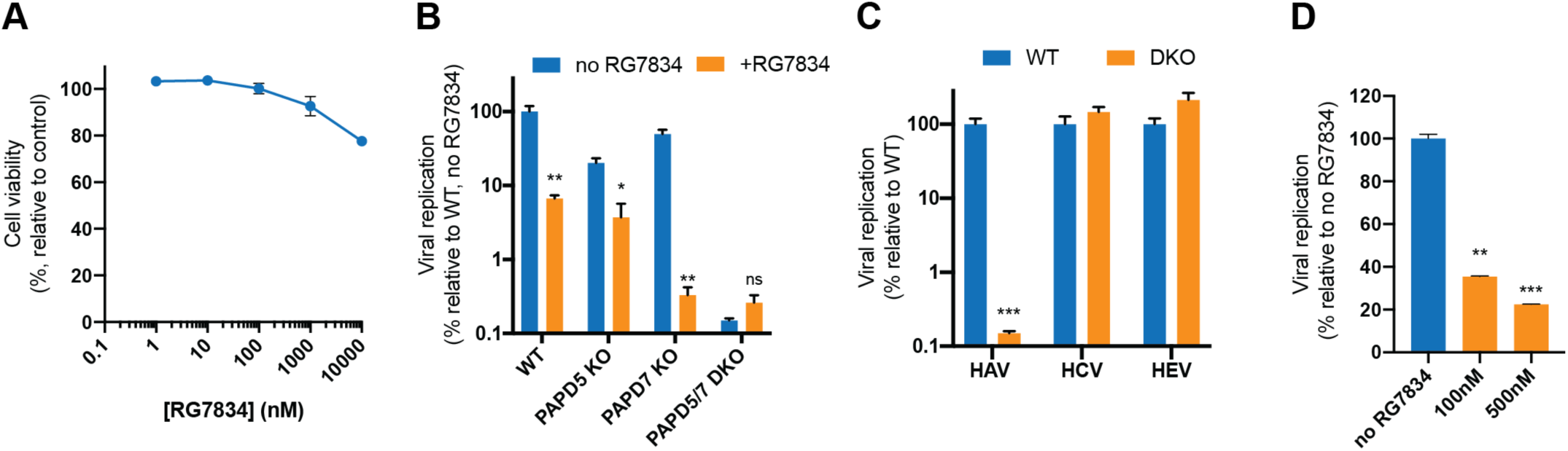
Small molecule inhibition of PAPD5 and PAPD7 reduces HAV infection in human hepatocyte cells and human liver organoids, Related to Figure 5. **(A)** Cell viability assay of Huh7.5.1 cells treated with different concentrations of RG7834 for 3 days. Dataset represents means ± s.d. (n=3 biological replicates) for each time point and values are relative to DMSO treated control. **(B)** Quantification of HAV RNA in WT, *PAPD5* KO, *PAPD7* KO or DKO cells untreated or treated with 100 nM RG7834 3 dpi by RT-qPCR. **(C)** Quantification of HAV and HCV RNA 3 dpi, and of HEV RNA 5 days post viral RNA transfection in WT or DKO cells using RT-qPCR. **(D)** Quantification of HAV RNA in human liver organoids treated with no, 100 nM or 500 nM RG7834 by RT-qPCR. For RT-qPCR experiments, viral RNA was normalized to 18S RNA and results are displayed relative to infection in (untreated) WT cells. Datasets represent means ± s.e.m. (n=3 biological replicates). P values were determined by unpaired two-sided t-tests using GraphPad Prism 8. ns, non-significant; *, p<0.05; **, p<0.01; ***, p<0.001; ****<0.0001.

## Methods

### Cell lines, viruses and reagents

Huh7.5.1 (gift from Frank Chisari), HEK293FT (Thermo Scientific) and U2OS cells (gift from Ron Kopito) were cultured in DMEM (Gibco) supplemented with 10% fetal bovine serum (FBS, Omega Scientific), penicillin/streptomycin (Gibco), non-essential amino acids (Gibco) and L-glutamine (Gibco) at 37C and 5% CO_2_. HepG2-NTCP-K7 cells (Ko et al., 2018) (gift from Ulrike Protzer) and HepAD38 cells (Ladner et al., 1997) (gift from Christoph Seeger) were cultured in DMEM/F12 (Gibco) supplemented with 10% FBS, penicillin/streptomycin, non-essential amino acids, L-glutamine and sodium pyruvate (Gibco) at 37C and 5% CO_2_. Cell lines were tested negative for mycoplasma and were authenticated by STR profiling. Hepatitis A virus HM175/18f (NR-137) was obtained through BEI Resources (NIAID, NIH) and propagated in Huh7.5.1 cells grown in adeno expression media (Gibco) supplemented with penicillin/streptomycin, non-essential amino acids and L-glutamine. Supernatant was collected 5-12 dpi and filtered (0.45 micron). To generate concentrated HAV stocks, supernatant was incubated with 8% PEG-8000 at 4C overnight, centrifuged at 3,200g for 60min and pellet was resuspended in a small volume of DMEM with 10% FBS and aliquoted. For the generation of HAV-Nluc (gift from Stanley Lemon) (Rivera-Serrano et al., 2019), the infectious clone plasmid was linearized, RNA was *in-vitro* transcribed using MEGAscript T7 Transcription Kit (Invitrogen) and transfected into Huh7.5.1 cells using TransIT-LT1 Transfection Reagent (Mirus Bio). Subsequently, cell supernatant was collected 8-13dpi, filtered and aliquoted. Purified encephalomyocarditis virus (NR-46441) was obtained through BEI Resources (NIAID, NIH). Enterovirus 71 (MP4), rhinovirus A2 and HCV JFH1 stocks were kindly provided by Jan Carette. For HBV production, supernatant was collected from HepAD38 cells, filtered (0.45 micron), and concentrated using 8% PEG-8000 as described above. The HEV infectious clone (genotype 3) was obtained from the NIH/NIAID and viral RNA was generated by *in-vitro* transcription from linearized plasmid (Shukla et al., 2012). For virus stocks, titers were determined by plaque-or focus-forming assay. RG7834 was purchased from MedKoo Biosciences and resuspended in DMSO at 50mg/ml. Puromycin and blasticidin were obtained from Gibco.

### CRISPR host factor screen

Huh7.5.1 cells were stably transduced with lentivirus from lentiCas9-Blast (Addgene #52962, gift from Feng Zhang) and subsequently selected using blasticidin. Next, a total of 300 million Huh7.5.1-Cas9 cells were then separately transduced with the lentiviral gRNA sublibraries A and B of the human GeCKO v2 library (Addgene #1000000049, gift from Feng Zhang) at a multiplicity of infection (MOI) of 0.3. Each sublibrary contained 3 guides per gene. Subsequently, the cells were selected using puromycin and expanded for 10 days. A total of 60 million mutagenized cells for each sublibrary (A and B) were collected for genomic DNA extraction, and a total of 45 million mutagenized cells for each sublibrary were infected with 15ml of unconcentrated HAV HM175/18f per T175 flask (1:1 diluted with fresh DMEM media). An additional 15ml of media was added after 2h. Media was refreshed 4 and 8dpi and surviving cells were collected 12dpi for genomic DNA extraction using QIAamp DNA Blood Maxi Kit (Qiagen). gRNA encoding DNA sequences were amplified in a two-step nested PCR using KAPA HiFi HotStart ReadyMixPCR Kit (Kapa Biosystems) and sequenced on an Illumina NextSeq (High Output) (for primer sequences, see Table S3). For analysis, reads were aligned to gRNA reference table and enrichment of each gRNA was calculated by comparing the relative abundance in the selected and unselected cell population. Gene enrichment analysis was performed using the RIGER weighted sum algorithm as well as MaGeCK analysis (Li et al., 2014; Luo et al., 2008). Gene ontology analysis of enriched genes in target list (containing 72 genes with RIGER p-value < 0.001) was performed using GOrilla (Eden et al., 2009). For comparison with other CRISPR host factor screens, RIGER-based gene enrichment analysis was used for HCV and HBV screens, and MaGeCK-based analysis for EV-D68 and HRV-C15 screens.

### Generation of clonal KO cell lines

Oligos containing gRNA sequence were annealed and ligated into pX458 (Addgene #48138, gift from Feng Zhang). Cells were transfected with pX458 constructs using Lipofectamine 3000 (Invitrogen) and two days later GFP positive cells were single-cell sorted into 96-well plates using a Sony SH800 cell sorter. For genotyping, genomic DNA was isolated from obtained clones using QuickExtract (Lucigen), the gRNA-targeted sites PCR amplified and the products Sanger-sequenced. Obtained sequences were compared to reference sequences for indel mutations or large deletion. A list of all used gRNA oligo and genotyping primer sequences can be found in Table S3.

### Intracellular RNA RT-qPCR assays

Cells were plated in 96-well plates (in triplicates for each condition) and infected the next day with virus at the following MOIs: HAV (∼5-10 FFU/cell), RV-A2 (∼1 PFU/cell), EV-71 (∼0.2 PFU/cell), EMCV (∼0.2 PFU/cell), HCV (∼1 FFU/cell) and HBV (∼1 FFU/cell). Cells were harvested, lysates reverse transcribed and quantitative PCR performed on a Bio-Rad CFX96 Touch system using the Power SYBR Cells-to-CT kit (Invitrogen). All viral RNA levels were normalized to 18S levels. Primer sequences can be found in Table S3.

### Quantification of extracellular viral RNA

Cells were plated in 96-well plates (duplicates for each condition) and infected with HAV at an MOI of 5. After 4h, inoculum was removed and cells were washed with PBS. At 3 dpi 100μl of supernatant was collected and RNA was isolated using the Qiagen Viral RNA Mini kit. Eluted RNA was reversed transcribed and analyzed by qPCR as described above. For determination of genome copies in-vitro transcribed HAV RNA was serially diluted and also quantified by RT-qPCR.

### Viral attachment and entry assay

96-well plates containing WT, *SLC35A1* KO or *HAVCR1* KO Huh7.5.1 cells were chilled on ice before infection with HAV HM175/18f at an MOI of 500 FFU/cell. Cells were incubated for 1h on ice before moving them to a 37C incubator. At each harvest timepoint, cells were washed three times with PBS and processed for RT-qPCR as described above.

### HAV nanoluciferase reporter assay

Cells were plated in 96-well plates and infected the next day with equal amounts of HAV-Nluc. Lysates were harvested by washing once with PBS wash followed by addition of 22µl of Passive Lysis Buffer (Promega) under shaking for 15min. For luminescence readout, 15µl of lysate were mixed with 50µl of Nano-Glo assay buffer (Promega) in a flat bottom white-walled luminescence plate, incubated for 5min at room temperature and read on an EnVision plate reader (PerkinElmer).

### Complementation of KO cells with cDNA

UBA5 cDNA was obtained from Dharmacon (DNA Accession: BC009737, Clone ID: 3879061). To generate cDNA encoding catalytically inactive enzyme, a Cys250Arg mutation was introduced by PCR. Additionally, a C-terminal FLAG-tag was added to WT and CAT cDNA sequences. UFM1 was obtained from Dharmacon (cDNA Accession: BC005193, Clone ID: 3829206). N-terminal FLAG and C-terminal mutations (Gly83Ala or ΔC3 (deletion of GlySerCys)) were introduced by PCR. For RPL26, cDNA was obtained from OriGene (RC209922) and a C-terminal FLAG-tag was added. To introduce 5KR mutations, a reverse primer containing point 5 mutations was used. ZCCHC14 cDNA was obtained from Dharmacon (cDNA Accession: BC101478, Clone ID: 8068984). The N-terminus was extended using a Gblock (Integrated DNA Technologies) resembling the protein coding sequence for a longer isoform (1086 amino acids, ENST00000671377.1) to match the molecular weight of ZCCHC14 detected in WT Huh7.5.1 by immunoblot. PCR products were gel-purified and cloned into EcoRV-cut pLenti CMV Puro DEST (w118-1) (Addgene #17452, gift from Eric Campeau & Paul Kaufman) using NEBuilder HiFi DNA Assembly Master Mix (New England BioLabs). Lentivirus was produced in HEK293FT and collected 48h post-transfection. KO cells were transduced with filtered, lentiviral containing supernatant under addition of polybrene and selected using 4 μg/ml puromycin for 3-4 days. For PAPD5 and PAPD7, pCI3 plasmids with WT or catalytic mutant cDNA sequences were kindly provided by V. Narry Kim and were described previously (Lim et al., 2018). For transient complementation, plasmids were transfected into WT or DKO cells using TransIT-LT1 Transfection Reagent (Mirus Bio), and transfected cells were plated in 96-wells the next day for HAV infection. To generate stable cell lines, cDNA sequences were cloned into the PB-CMV-MCS-EF1α-Puro vector (PB510B-1, SBI System Biosciences) where the CMV promoter was replaced by a PGK promoter. Plasmids were co-transfected together with a plasmid expressing piggyBac transposase into *PAPD5* KO cells, which were subsequently selected with puromycin for 5 days starting 3 days post-transfection. All primer sequences can be found in Table S3.

### Immunoblotting

Cell were lysed using Laemmli SDS sample buffer containing 5% beta-mercaptoethanol and boiled for 10min. Lysates were separated by SDS-PAGE on pre-cast Bio-Rad 4-15% poly-acrylamide gels in Bio-Rad Mini-Protean electrophoresis system. Proteins were transferred onto PVDF membranes using Bio-Rad Trans-Blot Turbo transfer system. PVDF membranes were blocked with PBS buffer containing 0.1% Tween-20 and 5% non-fat milk. Blocked membranes were incubated with primary antibody diluted in blocking buffer and incubated overnight at 4C on a shaker. Primary antibodies were detected by incubating membranes with 1:4000 dilution of HRP-conjugated (Southern Biotech) or 1:5000 dilution of IRDye-conjugated (LI-COR) secondary anti-mouse and anti-rabbit antibodies for 1 h at room temperature. Blots were visualized using a ChemiDoc MP Imaging System (Bio-Rad). The following primary antibodies (and their dilutions) were used in this study: p84 (Genetex, GTX70220) at 1:1000, GAPDH (SCBT, sc-32233) at 1:1000, vinculin (SCBT, sc-73614), 1:1000, eIF4B (SCBT, sc-376062) at 1:250, VPS4A (SCBT, sc-393428) at 1:300, SLC35A1 (Proteintech, 16342-1-AP) at 1:250, HAV VP1 (LS Bio, LS-C137674-100) at 1:1000, FLAG M2 (Sigma, F1804) at 1:1000, UFM1 (Abcam, ab109305) at 1:1000, UBA5 (Bethyl, A304-115A) at 1:2500, UFSP2 (SCBT, sc-376084) at 1:100, RPL26 (Bethyl, A300-686A) at 1:1000, PAPD5 (Atlas Antibodies, HPA042968) at 1:1000 and ZCCHC14 (Bethyl, A303-096A) at 1:1000. For SLC35A1, lysates from 10^6^ cells were immunoprecipitated using 4 µg anti-SLC35A1 antibody and Protein G magnetic Dynabeads (Thermo Scientific, 10-003-D) overnight at 4C prior to immunoblotting to enhance signal and specificity of signal.

### Replicon assay

The replicon plasmids pLuc-HAV/18f and pLuc-HAV/18f-3D^pol^GDD-->GAA (replication-defective mutant) were kindly provided by Stanley Lemon and were described previously (González-López et al., 2018). Plasmids were linearized using MluI-HF (New England BioLabs), RNA was generated using the MEGAscript T7 Kit (Invitrogen) and subsequently purified by lithium chloride precipitation. For electroporation, 1-2 million cells were washed three times in PBS, resuspended in 100μl SF Nucleofector solution (Lonza), mixed with 250ng replicon RNA per 80k cells, transferred to a 100 μl nucleocuvette and pulsed using the program FF-137 on an Amaxa 4D-Nucleofector X Unit (Lonza). Cells were then resuspended in equilibrated, antibiotic-free medium, distributed into 96-wells and lysed at different timepoints post-electroporation using 40μl Passive Lysis buffer (Promega). Luminescence was measured using Luciferase Assay System (Promega) on a white-walled luminescence plate with an EnVision plate reader (PerkinElmer).

### Poly(A) tail length assay by polyacrylamide gel electrophoresis

Cells were infected with HAV HM175/18f (MOI=10) and harvested 3 days post-infection using Trizol (Invitrogen). RNA was purified with Direct-zol RNA Microprep columns (Zymo Research). HAV poly(A) tails were analyzed using the Poly(A) Tail-Length Assay Kit (Thermo Scientific) according to the manufacturer’s instructions. Briefly, 1μg total RNA per condition was used as input for G/I tailing reaction followed by reverse transcription. Next, RT products were amplified using either the genome-specific (GS) forward and GS reverse primer, or GS forward and G/I specific reverse primer. PCR products were analyzed using polyacrylamide gel electrophoresis on a 5% Mini-PROTEAN Tris/Boric Acid/EDTA (TBE) gel in TBE buffer (100V for 60min). Gel was stained with SYBR Gold (diluted 1:10000 in TBE buffer) for ∼30min in the dark and imaged on a ChemiDoc MP Imaging System (Bio-Rad).

### Poly(A) tail sequencing

RNA was isolated from infected cells as described above. Ribosomal RNA (rRNA) was depleted from 1μg total RNA by addition of rRNA Removal Mix – Gold (Illumina) followed by incubation at 68C for 5min, addition of rRNA Removal Beads (Illumina), removal of supernatant while tubes are on a magnetic rack, and finally elution of rRNA-depleted RNA. Next, a 5’ adenylated, 3’ blocked oligodeoxynucleotide RNA linker (S1315S, New England BioLabs) was ligated to the 3’ ends of RNAs by incubation with RNA ligase 2, truncated (New England BioLabs) for 3h at 22C. Ligation products were then reverse transcribed using a primer, which is reverse complement to the RNA linker, and Superscript IV (Invitrogen). Finally, HAV poly(A) tails were specifically amplified in a nested PCR, cloned into the pCRBlunt II-TOPO vector (Invitrogen) and Sanger sequenced (Quintarabio). Number of adenosines downstream of the HAV 3’UTR region and occurrence of non-adenosine nucleotides were counted.

### Tapestation analysis of poly(A) tails of replicon RNA

10^6^ cells were electroporated with *in-vitro* transcribed replicon RNA containing a 40bp long poly(A) tail as described above. For each timepoint, cells were washed with PBS, trypsinized, pelleted and lysed using Trizol followed by RNA extraction with Direct-zol columns. Total RNA was G/I tailed, reverse transcribed and poly(A) tail region was amplified using the GS forward primer and G/I reverse primer. Products were analyzed using a High Sensitivity D1000 ScreenTape System (Agilent).

### Cell viability assay

Huh7.5.1 cells were treated with different concentrations of RG7834 for 3 days and viability was measured using Cell Titer Glo (Promega) according to the manufacturer’s instructions.

### Culture and HAV infection of human liver organoids

Bi-potent stem cell organoids were generated as previously described (Broutier et al., 2016; Huch et al., 2015). Briefly, liver tissue samples from healthy resection margins of partial hepatectomies were obtained through the Ibrahim El-Hefni Liver Biorepository at the California Pacific Medical Center Research Institute. Following tissue digest, the heterogeneous mixture of single cells was suspended in reduced growth factor BME2 (Basement Membrane Extract, Type 2, Trevigen). From this mixture, 50 µL drops containing 1,000 to 20,000 cells were seeded in 24-well suspension culture plates (GreinerBio). Drops were incubated at 37C for >20 min and solidified. After this, 500 µL of expansion media (EM) was added to each well. Expansion media is basal media (Advanced DMEM/F12 with 1% penicillin/streptomycin, 1% Glutamax, and 10 mM HEPES (all from ThermoFisher) supplemented with 1% N2 (Gibco), 1% B27 (Gibco), 1 mM N-Acetylcysteine (Sigma-Aldrich), 10 nM [Leu^15^]-gastrin I human (Sigma-Aldrich), 10% (vol/vol) R-spondin1 conditioned media (generated as described in (Broutier et al., 2016)), and 10 mM Nicotinamide (Sigma-Aldrich). Expansion media additionally contains 50 ng/ml recombinant human EGF, 25 ng/ml recombinant human HGF, 100 ng/ml recombinant human FGF10, 10 µM Forskolin, and 5 µM A83-01 (all from Stem Cell Technologies). Expansion media was replaced every 3-4 days. Organoids in expansion media grew in culture for 4-6 months and needed to be routinely passaged with TRYPLE (Gibco). To induce differentiation to a hepatocyte-like fate, expansion media was supplemented with 25 ng/ml BMP7 (ProSpec) for 3-4 days. After this, media was changed to differentiation media (DM). Differentiation media is basal media supplemented with 1% N2, 1% B27, 1 mM N-acetylcysteine, 10 nM [Leu^15^]-gastrin I human, 50 ng/ml EGF, 25 ng/ml HGF, 0.5 µM A83-01, 25 ng/ml BMP7, 3 µM dexamethasone (Sigma-Aldrich), 10 µM DAPT (Stem Cell Technologies), and 100 ng/ml recombinant human FGF19 (ProSpec). Differentiation media was changed every 3-4 days for a period of 3-15 days. For HAV infection, organoid lines from two non-viral donors were spin-infected with the HM175/18f strain of HAV as follows. Organoids from EM and DM d3 conditions were collected and lightly dissociated by a 3 min incubation at 37°C with TRYPLE. Cells were then suspended in basal media containing 10 µM Y-27632 (Stem Cell Technologies) and counted. These cells were mixed with HAV at an MOI of 500 FFU/cell in the presence and absence of RG7834 at 100 or 500 nM. Cell suspensions were then added to a 24-well suspension culture plate. The plate was centrifuged at 600 x g for 1 h at room temperature, followed by a 2 h incubation at 37C. After this, cells were collected and washed 3x in basal media. Washed cells were seeded in fresh BME2 drops in a new 24-well suspension culture plate. At 7 days post-infection, infected organoid samples were washed 2x in cold basal media, and cell pellets were lysed in Trizol.

### Statistical analysis

Prism 8 (GraphPad Software) was used for statistical analysis. Statistical tests and p-value definitions are listed in the figure legends.

## Supplementary files

Table S1: Gene enrichment analysis of HAV CRISPR screen. Table S2: Gene ontology analysis for hits of HAV CRISPR screen. Table S3: Oligo sequences.

## Data availability

Raw sequencing data for CRISPR KO screen is deposited at the EMBL-EBI ArrayExpress (https://www.ebi.ac.uk/arrayexpress/) under the accession number E-MTAB-8646.

## Author contributions

JK and ASP were responsible for design and execution of experiments, data analysis and manuscript preparation. RW performed comparative analysis of CRISPR screens and assisted with molecular biology experiments. NLM conducted organoid experiments under the supervision of MO. ASP conceptualized and supervised the study.

## Acknowledgments

We would like to thank Don Ganem for helpful discussions and critically reading the manuscript, Sandra Schmid for valuable feedback on the manuscript, the Biohub Genomics platform for help with sequencing, James Webber for assistance with the CRISPR screen analysis, and members of the Biohub Infectious Disease Initiative for helpful discussions. We would like to acknowledge Drs. Stanley Lemon (University of North Carolina, Chapel Hill), Jan Carette (Stanford University), V. Narry Kim (Seoul National University), Ron Kopito (Stanford University), Ulrike Protzer (Technical University Munich), Feng Zhang (Broad Institute), Christoph Seeger (Fox Chase Cancer Center) and Suzanne Emerson (NIH) for providing critical reagents. We also acknowledge Drs. Stewart Cooper and Ann Erickson, who oversee the Ibrahim El-Hefni Liver Biorepository at the California Pacific Medical Center Research Institute and provided patient tissue samples used to generate liver organoids. The research was funded by the Chan Zuckerberg Biohub (ASP) and by NIH/NIDA grants AI097552 and DP1DA038043-01 (MO).

